# Biolog phenotype microarray: a tool for the identification of multidrug resistance efflux pumps inducers

**DOI:** 10.1101/344879

**Authors:** P. Blanco, F. Corona, JL. Martínez

## Abstract

Overexpression of multidrug resistance efflux pumps is a relevant mechanism of antibiotic resistance for bacterial pathogens. These systems use to present low levels of basal expression. However, they can be induced by environmental signals or stresses which can lead to situations of phenotypic induced resistance. In contrast to efflux pumps substrates, inducers of these systems have not been thoroughly studied. In this work, we have applied a novel high-throughput methodology in order to identify inducer molecules of the *Stenotrophomonas maltophilia* SmeVWX and SmeYZ efflux pumps. To that goal, bioreporters in which the expression of the yellow fluorescent protein is linked to the activity of either the *smeVWX* or the *smeYZ* promoters were developed and used for the screening of potential inducers of the expression of these efflux pumps using Biolog phenotype microarrays. Confirmation of induction was carried out measuring YFP production along the bacterial growth and by flow cytometry; mRNA levels of *smeV* and *smeY* were also determined by real-time RT-PCR after exposure to the selected compounds. Among the 144 tested compounds, iodoacetate, clioquinol (5-chloro-7-iodo-8-hydroxyquinoline) and sodium selenite were found to be *smeVWX* inducers, while boric acid, erythromycin, chloramphenicol and lincomycin are able to trigger the expression of *smeYZ*. While the presence of the inducers allowed a decrease in the susceptibility to antibiotics that are known substrates of the efflux pumps, our results indicate that these efflux pumps did not contribute to *S. maltophilia* resistance to the analyzed inducers.

**Importance:** Multidrug efflux pumps constitute a category of elements involved in the cellular response to stress that is universally represented; from bacteria to human cells. Besides playing basic roles in cell physiology, these elements are critical elements in the resistance to therapeutic agents, including anti-cancer drugs, antifungals and antibiotics. Stable-inheritable resistance is achieved through mutations in regulatory elements that allow overexpression of these systems. However, much less is known on the effectors, or growing conditions, that might induce their expression, leading to a situation of transient-phenotypic resistance, not detectable by current susceptibility tests, unless the inducer in known. Herein we present a methodology amenable for the high-throughput screening of efflux pumps inducers. The use of phenotype microarrays linked to fluorescence reporters have allowed to identify a set of different inducers for *smeVWX* and *smeYZ*. Notably, induction seems to be uncoupled from the detoxification of the inducers by the corresponding efflux pumps. The mechanism of action of each of the inducers for inhibiting bacterial growth allowed us to propose that *smeVWX* is likely induced as a response to thiol-reactive compounds, while *smeYZ* is induced by ribosome-targeting antimicrobials. Although applied to a specific bacterium, this method is of application to any type of organism and efflux pump, changing the growing conditions in the case of eukaryotic cells. Since the presence of inducers may change the cell response to therapeutic drugs, the identification of these molecules is of clinical relevance.

## Introduction

Multidrug resistance (MDR) efflux pumps play a key role in antimicrobial resistance through the active transport of antimicrobial agents outside the cell. Depending on their basal level of expression, they can contribute to the intrinsic resistance to different compounds displayed by bacterial pathogens. In addition, the overexpression of MDR efflux pumps also leads to high levels of resistance to clinically useful antibiotics (1). The resistance nodulation division (RND) MDR family is the best characterized one. Members of this family are present just in Gram-negative bacteria, being associated with clinically relevant antimicrobial resistance (2, 3). While RND systems are usually expressed at very low level (if any) under regular growing conditions, a diversity of complex pathways is involved in the regulation of their expression, which may require the participation of several global and local transcriptional regulators belonging to different families, including TetR, LysR or two-component systems (TCS) (4).

*Stenotrophomonas maltophilia*, a relevant emerging MDR opportunistic pathogen, exhibits low susceptibility to a wide range of antibiotics due the presence of several resistance mechanisms encoded in its genome, including eight MDR efflux pumps belonging to the RND family (5-7). *S maltophilia* RND efflux pumps exhibit different patterns of regulation, which makes this organism a suitable model for the investigation of these systems. For instance, the expression of *smeYZ* efflux pump is linked to the TCS SmeRySy (8), and *smeVWX* expression is regulated by the LysR-type regulator SmeRv (9). In addition, whereas *smeYZ* is constitutively expressed at a significant level, hence contributing to intrinsic resistance to aminoglycosides (10, 11), *smeVWX* expression level is low and resistance is only achieved through mutations in its regulator, leading to the overexpression of the efflux pump (12).

It is important to recall that overexpression of RND efflux pumps is not always associated with genetic changes. Indeed, efflux pumps are expected to be expressed, when needed, in response to specific signals/cues (2). In this respect, some situations can lead bacteria to become transiently resistant, such as the presence of compounds that act as effectors, triggering the expression of these MDR systems and displaying a phenotype of transient resistance (13, 14). These situations depend on the regulatory network of the RND efflux pumps, including local and global regulators. Most of the RND effectors have been discovered through the study of specific physiological processes wherein efflux pumps participate, such as the colonization of particular niches, the extrusion of a particular toxic compound or the characterization of specific regulators (15-17). Besides the assigned specific function of RND efflux pumps, these systems are in occasions considered stress defense determinants (e.g. oxidative and nitrosative stress), acting as scape valves for the accumulation of toxic compounds or stress by-products (18).

RND efflux pumps are known to extrude a wide range of structurally different compounds, some of which are usually well-characterized, especially when they are clinically relevant antimicrobials (19). However, less is known about the efflux pump effectors. Knowing the set of RND inducer molecules could be helpful for characterizing MDR efflux systems, getting understanding on their potential functional and ecological roles, as well as for detecting possible situations of phenotypic induction of resistance *in vivo* that would not be detected by classical laboratory susceptibility tests. In this study, we have performed a systematic analysis of potential inducers of the expression of the efflux systems SmeVWX and SmeYZ applying the phenotype microarray technology, providing new insights about their regulation mechanisms and their role in the physiology of *S. maltophilia*.

Biolog phenotype microarrays consist in 96-well microplates developed for the determination of bacterial growth phenotypes under different conditions. These microplate assays are designed for measuring the response of a microorganism to a wide variety of chemical agents and nutrients, allowing the testing of nearly 2.000 phenotypes (20). The system is based on a colorimetric reaction using tetrazolium dye for detecting cellular respiration as an indicator of an active cellular metabolism (21). These phenotype microarrays can be used for different purposes, including the testing of biofilm formation (22), bacterial evolution experiments (23), studies on fitness costs associated to the acquisition of resistance to antimicrobials (24-26), or the characterization of knock-out mutants concerning their capacity to metabolize different carbon sources or their survival in the presence of toxic compounds (27-29). In this study, we have employed for the first time a different approach to perform a wide screening of efflux pumps’ inducer molecules, which could be applied as well to any other gene transcriptional regulation. Using fluorescent reporters of the RND efflux pumps SmeVWX and SmeYZ, we describe here the identification of a set inducers of these systems by measuring the fluorescence signals after the exposure to 144 compounds belonging to different categories. The global study has been completed with the analysis of the effect of induction on the susceptibility to the inducer compounds and to antibiotics currently used in clinical practice.

## Results

### Characterization of PBT03 (*P_EM7_*) and PBT10 (*P_YZ_*) reporter strains

With the aim of measuring the expression of *S. maltophilia* efflux pumps in response to potential effectors, the use of YFP-based reporters has been employed in this work. In a recent study, we developed the YFP-based reporter strain PBT02 in order to identify potential inducer compounds of the SmeVWX efflux pump in *S. maltophilia* (30). In the present work, two new reporter strains have been developed, one for measuring *smeYZ* expression, and another, which contains a constitutive promoter as a control for the normalization of the results. The promoter region of *smeYZ* (136 pb) was cloned into the pSEVA237Y plasmid, obtaining the pPBT11 plasmid, whereas the plasmid pPBT05 was obtained by cloning *EM7*, a synthetic bacterial promoter derived from the bacteriophage T7 (31), into the plasmid pSEVA237Y. Due to its high constitutive transcription activity, it can be used as control of expression. Both pPBT11 and pPBT05 plasmids were introduced in *S. maltophilia* D457 by tripartite conjugation generating PBT10 (*P_YZ_*) and PBT03 (*P_EM7_*) strains, respectively.

In order to test the proper functioning of both reporter strains, bacterial growth and YFP levels were measured for 20 h. Although the validation of PBT02 strain was performed in a previous work (30), it was also included in the analysis. The fluorescence levels given by the PBT10 strain are higher in comparison with those obtained for the already described PBT02 strain (Figure S1), confirming that *smeYZ* is constitutively expressed in the wild-type *S. maltophilia* strain (10), while *smeVWX* expression is very low. The YFP level given by the PBT03 strain, which harbors the plasmid containing the *EM7* promoter, were also assessed and, as expected, the expression level was high enough to be used as a positive control of fluorescence and for normalization purposes (Figure S1).

### Screening of inducers of MDR efflux pumps expression using Biolog phenotype plates

Four different quantities of each of the compounds are dried on the base of each well in the Biolog plates. We have denominated each well as A, B, C, and D, from the highest to the lowest concentration of the compound. For this study, plates PM11 to PM16, making a total of 144 analyzed compounds (Table S1), were chosen. *S. maltophilia* reporter strains grew in at least one concentration of 142 compounds (Figure S2). Those wells were *S. maltophilia* did not grow were discarded for further analysis.

Although Biolog microarray plates have been used for different purposes that do not involve the use of the tetrazolium dye, there are no reports of the employment of these plates in spectrofluorometric assays. The recorded fluorescence was analyzed in a two-step method (see Materials and Methods) to obtain a proxy value of the transcription activity. Briefly, the production of fluorescence was quantified for each curve giving rise to the maximum specific rate of fluorescence production parameter (i.e. *µ_EM7_ µ_YZ_* or *µ_VWX_* for each compound and concentration). Then, all values representing *P_YZ_* or *P_VWX_* transcription activity were normalized using the constitutive fluorescence given by the *P_EM7_* as control. This step is required in order to avoid any effect of the toxic compound on the fluorescence signal, which may be caused by itself or by its action on the growth rate, being inversely related to the fluorescence signal (32). Finally, the *µ_VWX_*/*µ_EM7_* and *µ_YZ_*/*µ_EM7_* values for each compound and concentration were used to represent the normalized transcription activity of *P_VWX_* and *P_YZ_*, respectively.

To stablish a threshold value that enables to define overexpression, all efflux pump transcription proxies’ values were depicted in a box plot (Figure 1). In this way, a threshold of *µ_VWX_*/*µ_EM7_*= 0.14 and *µ_YZ_*/*µ_EM7_* = 0.89 were set to define those situations where *smeVWX* and *smeYZ* were considered to be overexpressed. All the compounds and concentrations that gave rise to both efflux pumps overexpression are listed in Table 2. In the case of the SmeVWX efflux pump, 23 compounds and 34 concentrations were found to exceed the threshold, while 22 compounds and 39 concentrations were above the set threshold in the case of the SmeYZ efflux pump. For instance, sodium selenite and 5-chloro-7-iodo-8-hydroxyquinoline caused the overexpression of *smeVWX* in the four tested concentrations, and the same happens for *smeYZ* with protamine sulfate and 5-fluorootic acid. In the case of 18 compounds, only the maximum tested concentration led to the efflux pump overexpression.

**Figure 1.**
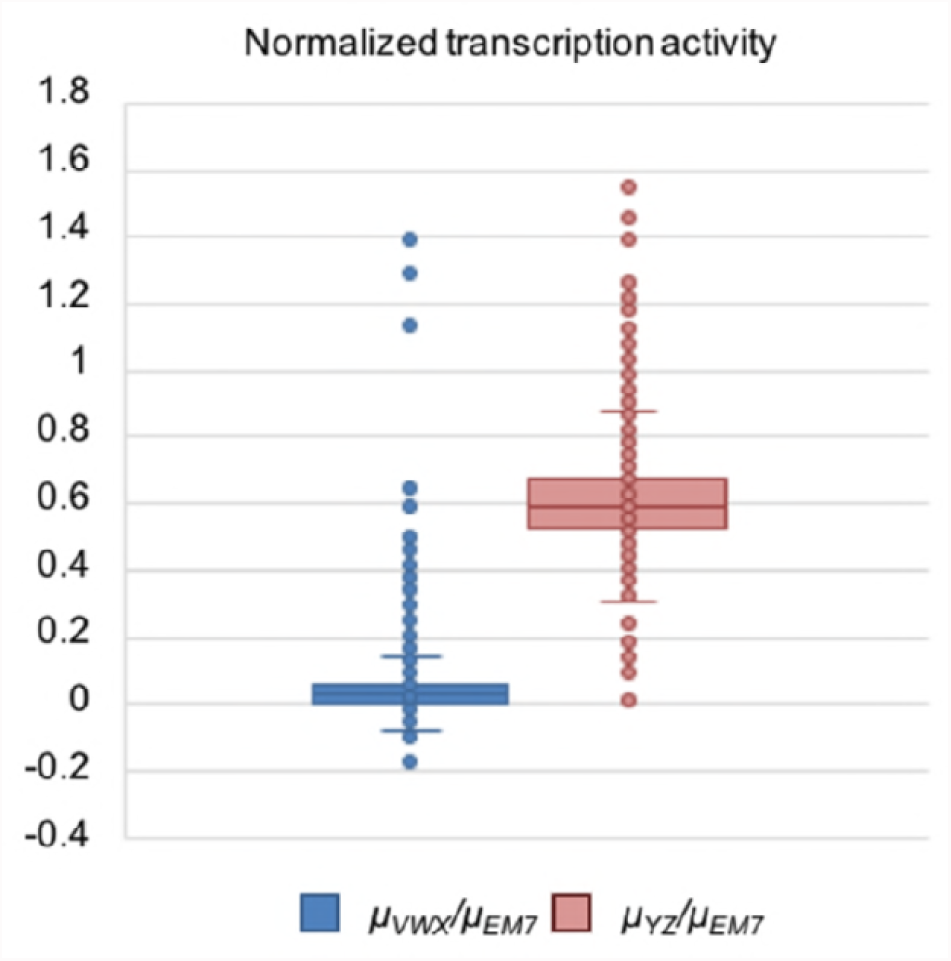
Normalized transcription activity. Box plot representing the values of *µ_RND_/µ_EM7_* for all Biolog compounds and concentrations tested in plates PM11-PM 16 where there was observable growth. The upper whisker delimits the threshold above which outlier transcription activity is defined for *smeVWX* or *smeYZ* promoters, and therefore efflux pump overexpression.

**Table 1.**
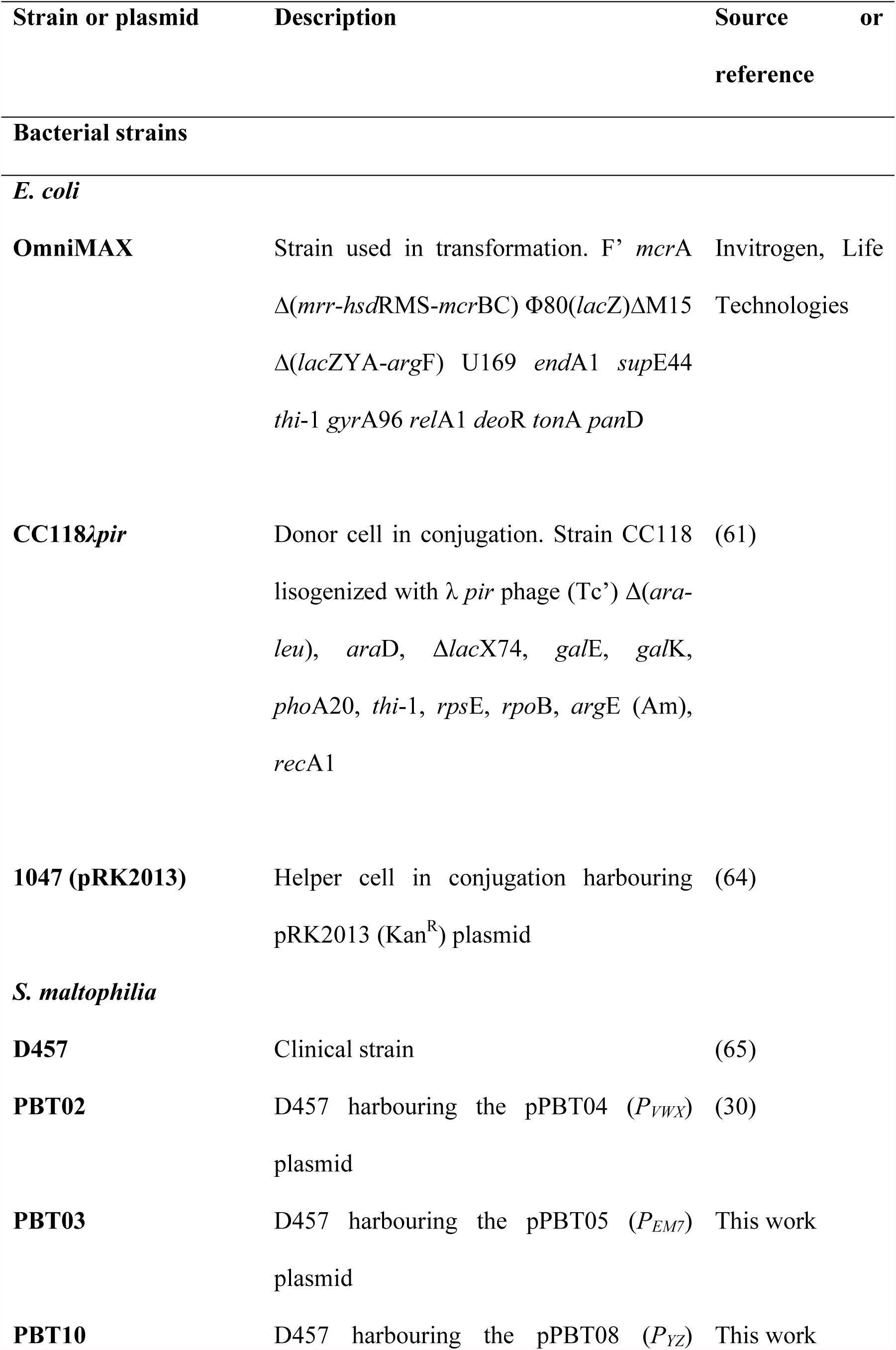
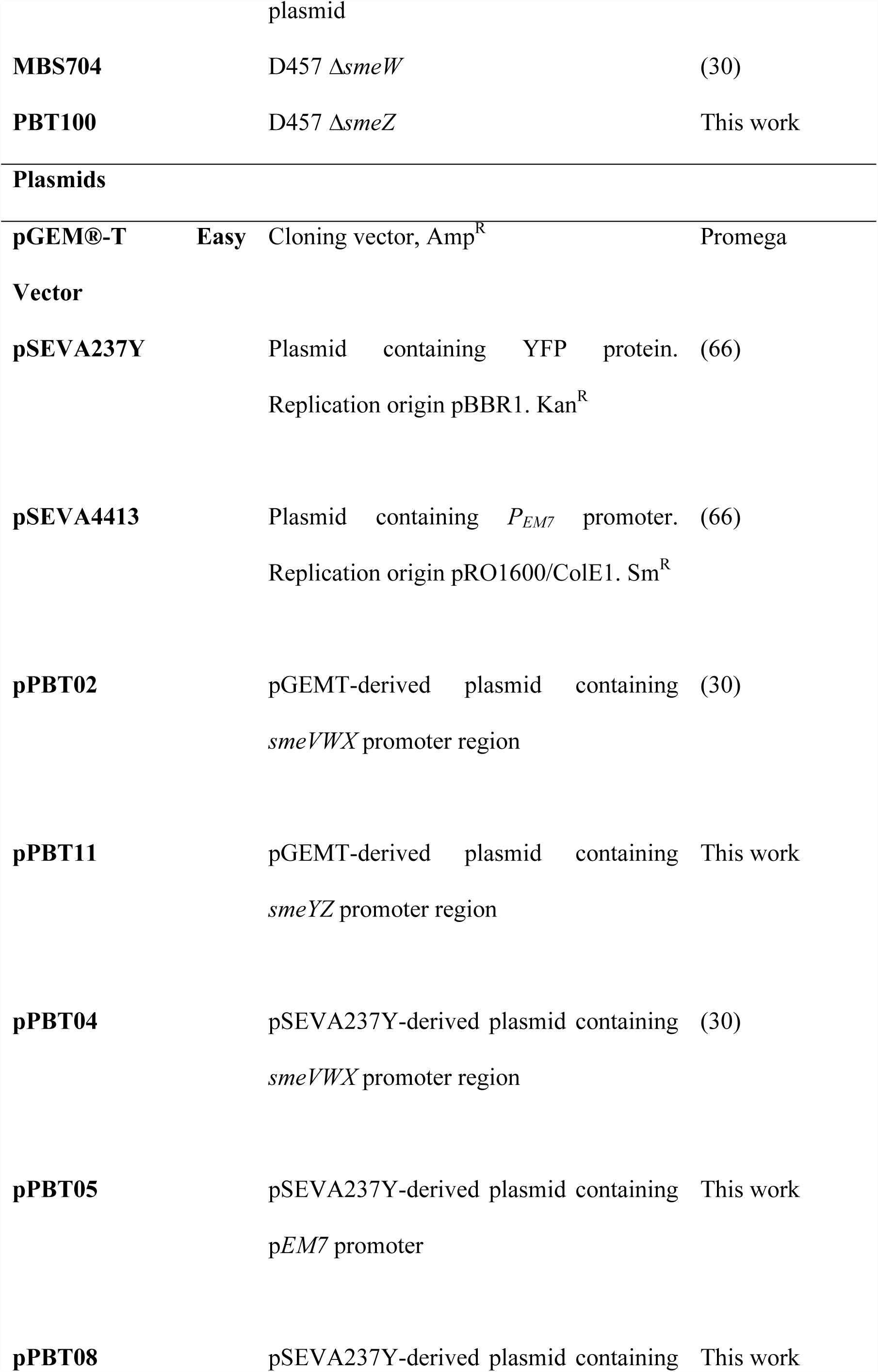
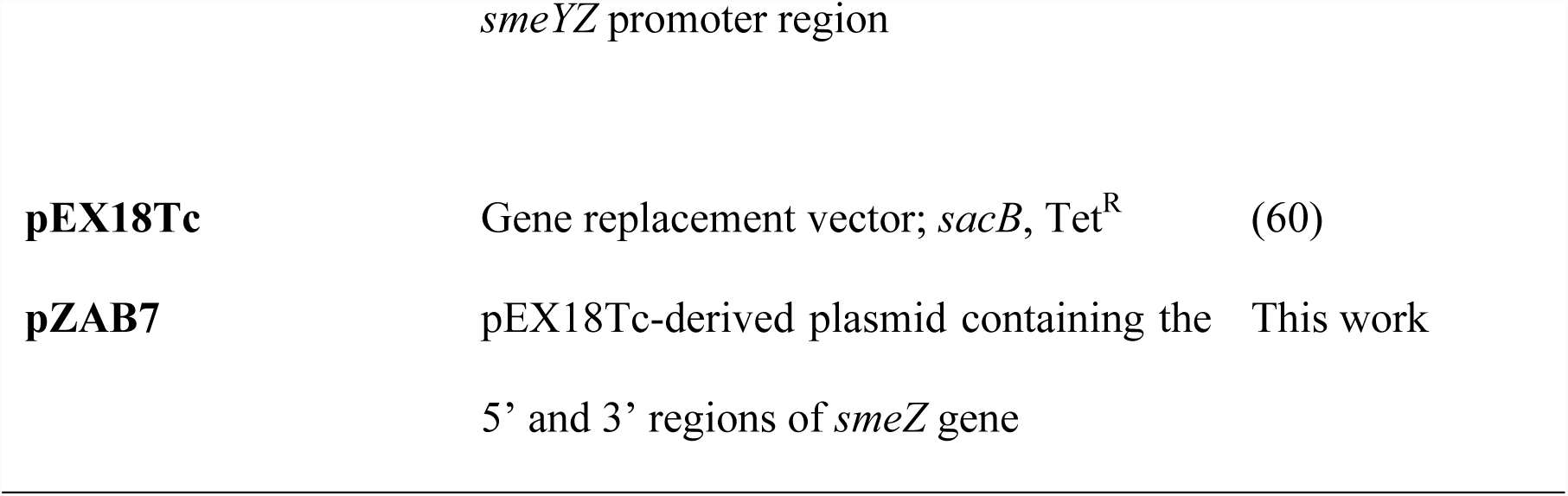
Bacterial strains and plasmids

**Table 2.**
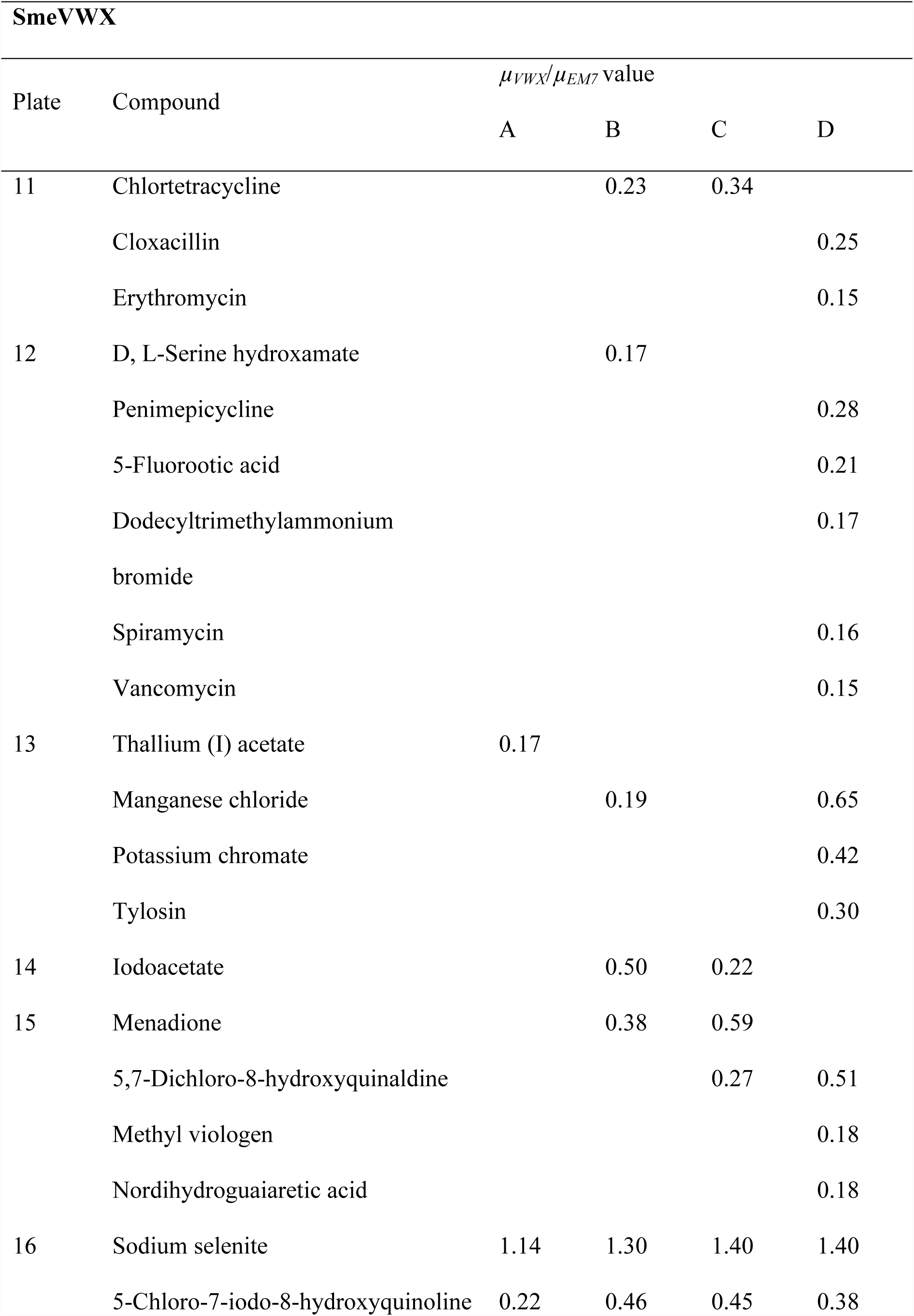
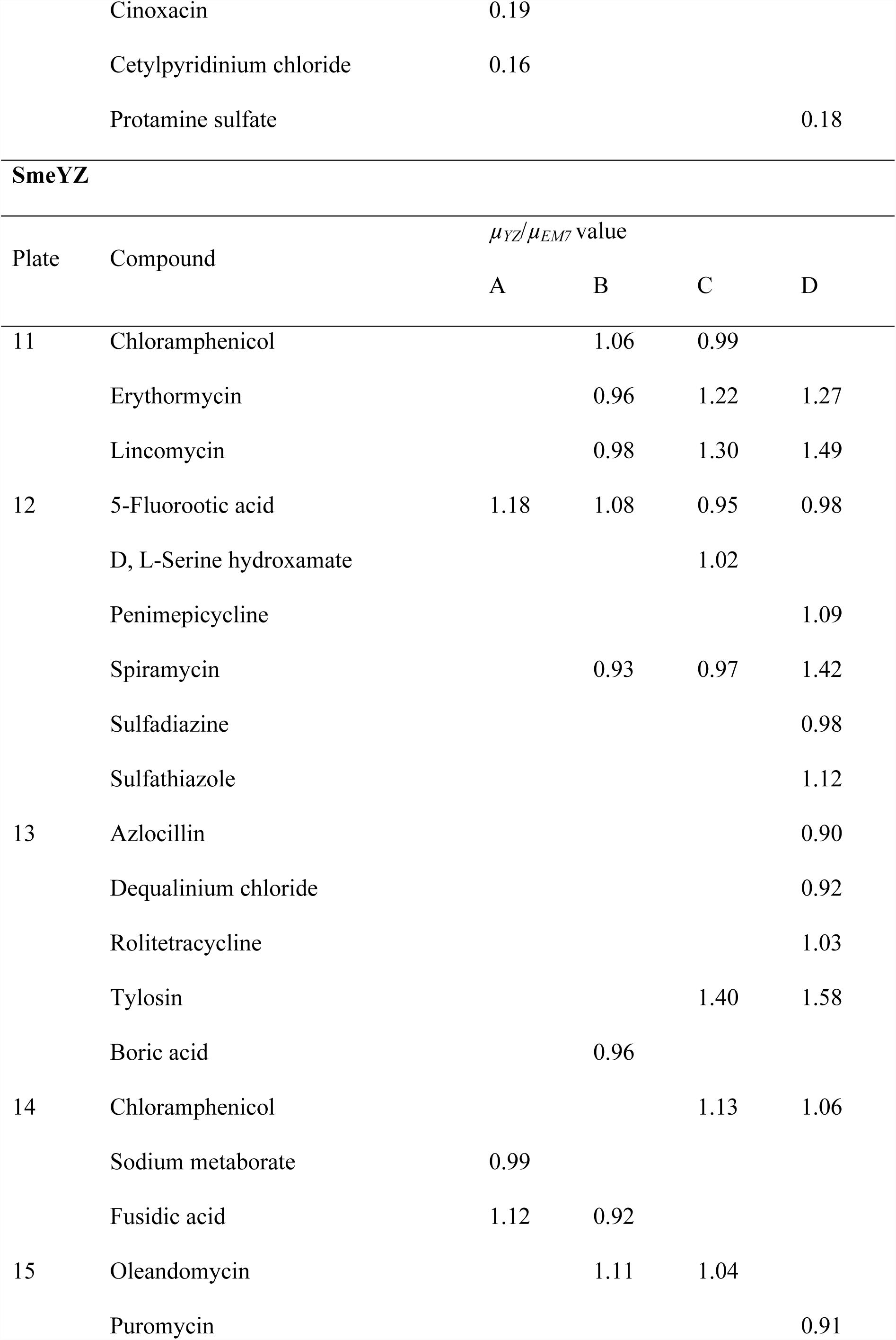
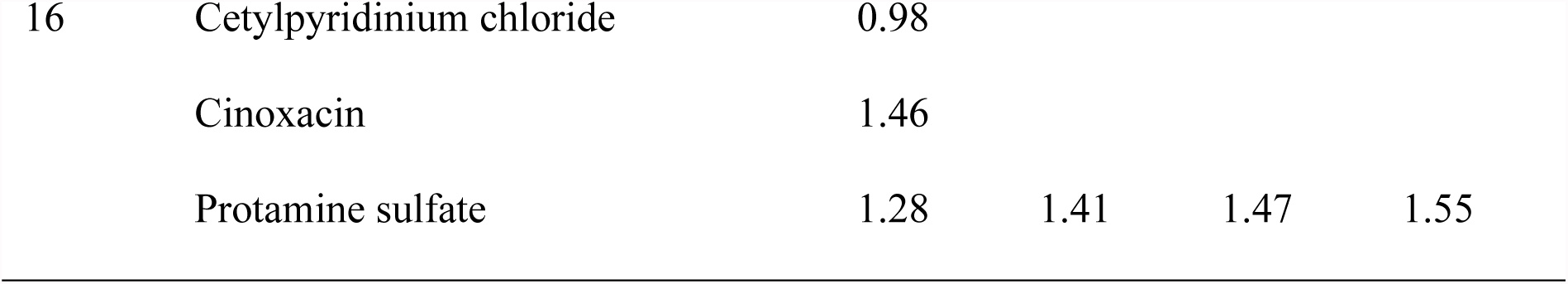
Normalized transcriptional activity of *P_VWx_* and *P_YZ_* with Biolog compounds

### Confirmation of the increased expression of *smeVWX* and *smeYZ* in the presence of potential inducers

Among those Biolog compounds that were able to produce an increase in the YFP levels for each of the studied efflux pumps, we selected some of those for which the induction was higher in order to validate the results. We selected as possible inducers of *smeVWX* iodoacetate, clioquinol (5-chloro-7-iodo-8-hydroxyquinoline) and sodium selenite; for potential *smeYZ* inducers, we selected boric acid, chloramphenicol, erythromycin and lincomycin. Since the quantity of each compound present in the Biolog plates is not available for users, we first determined the MIC in order to know the concentrations to be used in the following experiments. These concentrations were: iodoacetate, 1 mM; clioquinol, 0.065 mM; sodium selenite, ≥ 200 mM; boric acid, 25 mM; erythromycin, 256 µg/ml; chloramphenicol, 4 µg/ml; lincomycin, 4096 µg/ml. Then, we selected three concentrations below the MIC value for each compound in order to perform the induction assay.

To carry out the experiment, the reporter strains PBT02 (*P_VWX_*) and PBT10 (*P_YZ_*) were incubated with their respective putative inducers for 20 h. The PBT03 (*P_EM7_*) strain was incubated with all the compounds as a fluorescence control. In Fig. 2, the YFP values and growth given by the PBT02 and PBT10 strains are shown, using the concentration for which the highest induction was obtained and bacterial growth was less compromised in each case: 0.5 mM sodium selenite, 78 μM clioquinol, 0.5 mM iodoacetate, 6.25 mM boric acid, 8 μg/ml erythromycin, 1 μg/ml chloramphenicol, and 128 μg/ml lincomycin. In agreement with the Biolog data, iodoacetate, clioquinol and sodium selenite are able to increase the YFP production through the induction of the *smeVWX* promoter in the PBT02 strain, in comparison with the fluorescence observed in control medium without inducer (Figure 2A); while erythromycin, boric acid, chloramphenicol and lincomycin increase the YFP production through the induction of the *smeYZ* promoter in the PBT10 strain, when compared with the strain growing without these compounds (Figure 2B). PBT03 did not show any difference in YFP production when incubated with the different compounds or without them (Figure 2C). These results show that the observed fluorescence increment for both reporter strains PBT02 and PBT10 when growing in the presence of the tested putative inducers is due to the induction of the promoters of the tested efflux pumps and not an indirect effect of the compounds themselves.

**Figure 2.**
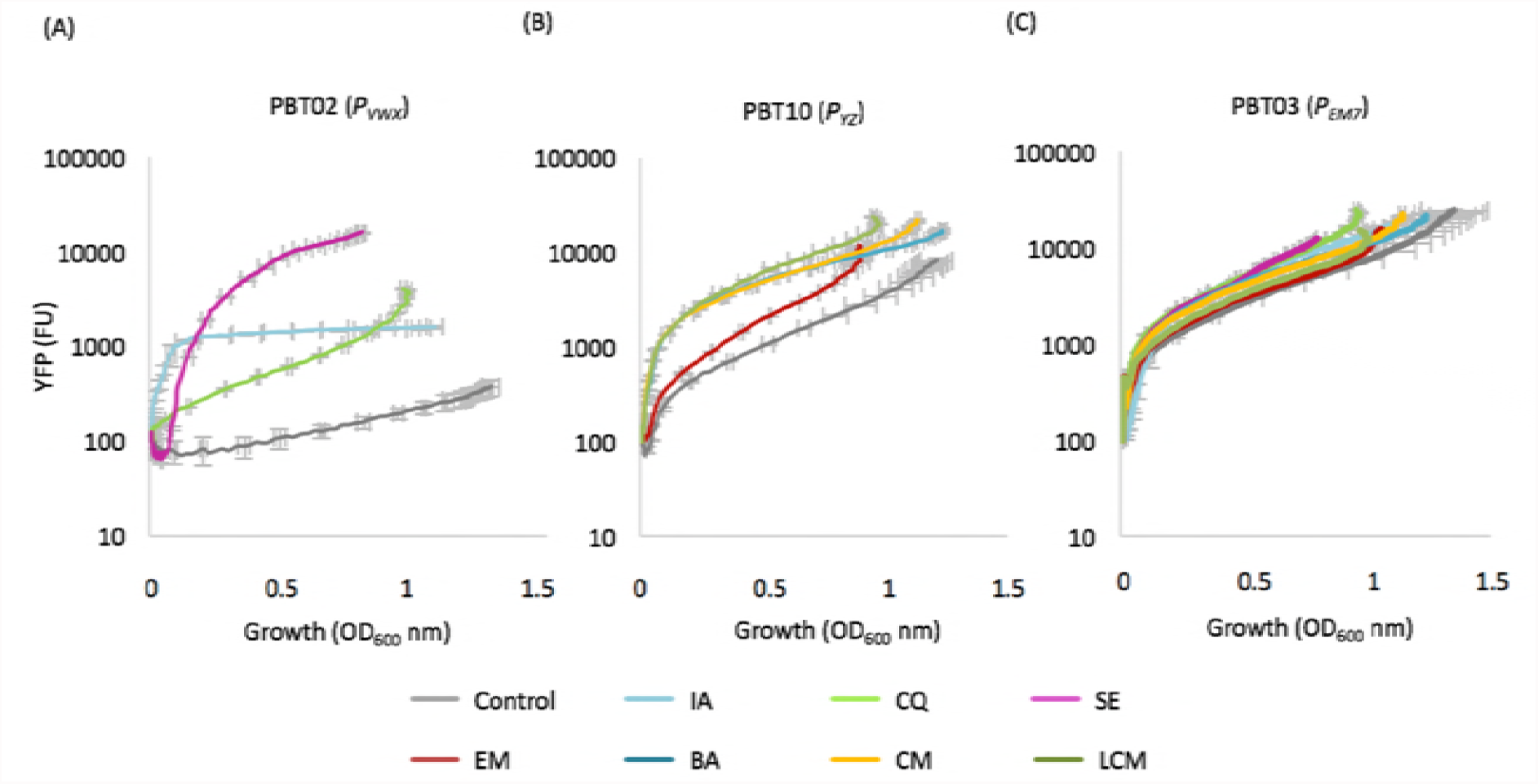
Growth and fluorescence values given by the reporter strains PBT02 (*P_VWx_*), PBT10 (*P_YZ_*) and PBT03 (*P_EM7_*) after incubation with the selected putative inducers. (A) Expression of *smeVWX* during 20 h incubation in the presence of iodoacetate (0.5 mM), clioquinol (78 µM) or sodium selenite (0.5 mM). (B) Expression of *smeYZ* during 20 h incubation in the presence of erythromycin (8 µg/ml), boric acid (6.25 mM), chloramphenicol (1 µg/ml) or lincomycin (128 µg/ml). (C) *EM7* expression during 20 h incubation with all the compounds at the above mentioned concentrartions. Error bars show standard deviations from three independent replicates. FU, fluorescence units. CQ, clioquinol; IA, iodoacetate; SE, sodium selenite; BA, boric acid; CM, chloramphenicol; EM, erythromycin; LCM, lincomycin.

### Single-cell analysis of the induction of the expression of MDR efflux pumps

In order to examine the effect of induction on the activity of *smeVWX* and *smeYZ* promoters at single-cell level, we measured the YFP values by flow cytometry given by the two reporter strains PBT02 and PBT10 after 90 min of incubation with their respective inducers when cells reached exponential growth phase (OD_600_ ≈ 0.6) (Figure 3). This methodology lessens any time-dependent interference with the fluorescence signal, such as the bacterial growth phase or compound degradation. PBT03 was also treated with all the set of compounds as a control (Figure 3C and 3D). As shown in Fig. 3A, the bacterial populations of PBT02 strain incubated with clioquinol, iodoacetate or sodium selenite present a higher *smeVWX* expression than the untreated population. The same results are obtained when the PBT10 strain is incubated with boric acid, erythromycin, chloramphenicol or lincomycin, showing these compounds to induce *smeYZ* expression (Figure 3B). For all the compounds, a normal distribution of the level of expression was seen in the population, indicating that all cells presented an increased expression of the corresponding efflux pump in presence of the inducers. In addition, the expression levels correspond with those obtained in the above growing curves, confirming that induction of efflux pumps promoters is produced at mid-exponential growth phase.

**Figure 3.**
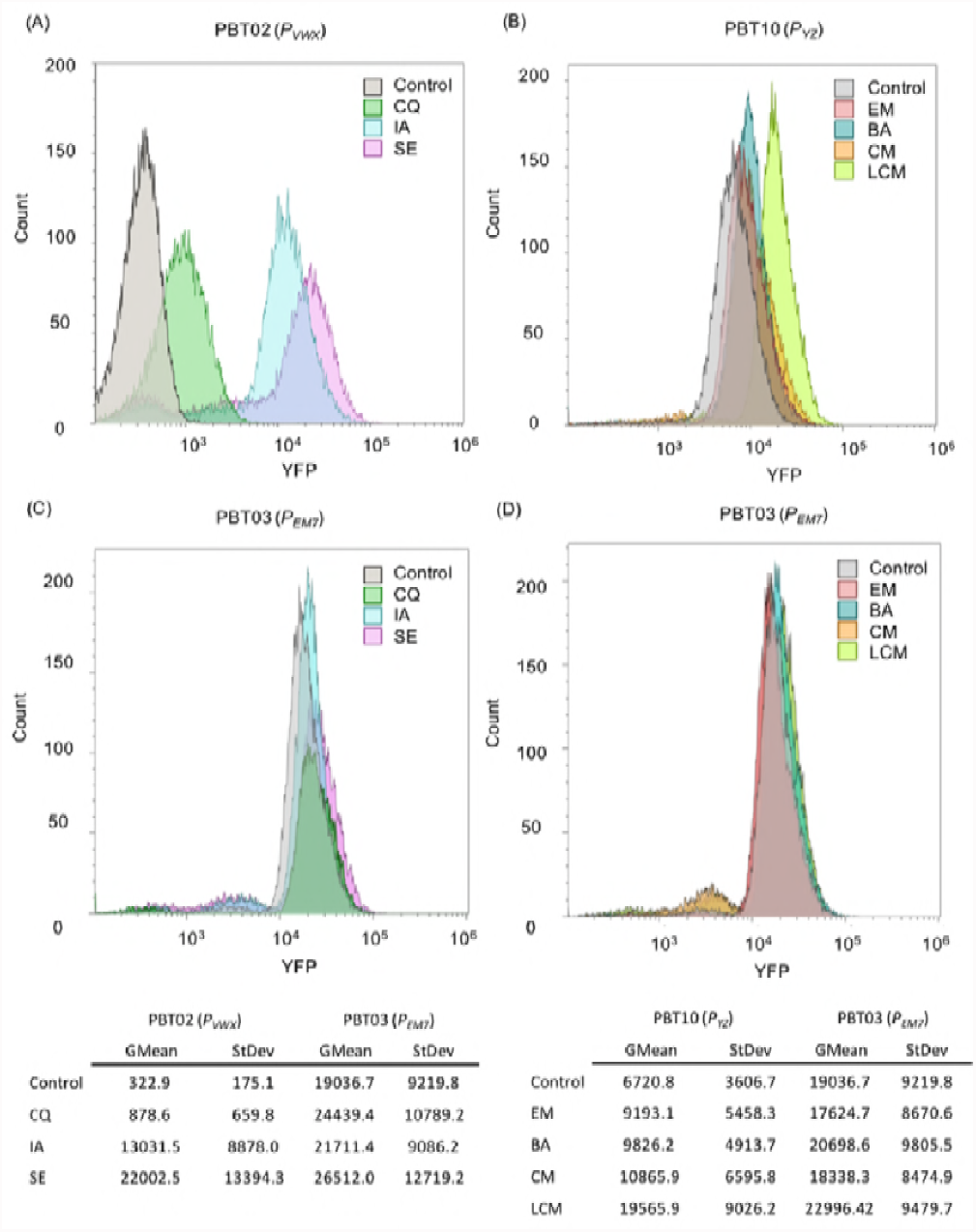
Population analysis of *smeVWX* and *smeYZ* expression in the presence of the selected putative inducers. YFP production was analyzed by flow cytometry. (A) Expression profile of *smeVWX* promoter in PBT02 strain treated with clioquinol (78 µM), iodoacetate (0.5 mM) or sodium selenite (0.5 mM). (B) Expression profile of *smeYZ* promoter in PBT10 strain treated with erythromycin (8 µg/ml), boric acid (6.25 mM), chloramphenicol (1 µg/ml) or lincomycin (128 µg/ml). (C) and (D) Expression profile of *EM7* promoter in PBT03 strain treated with all the different compounds at the same above mentioned concentrations. Gray populations represent in all cases the basal promoters’ expressions in the absence of any compound. Geometric mean (GMean) and standard deviation (StDev) were calculated for each population using Kaluza 1.5 software. CQ, clioquinol; IA, iodoacetate; SE, sodium selenite; BA, boric acid; CM, chloramphenicol; EM, erythromycin; LCM, lincomycin.

### Analysis of the expression of efflux pumps in the presence of inducers at mRNA level

For further confirmation of the results, the mRNA levels of *smeV* and *smeY* were quantified by real-time RT-PCR in the presence of the different inducer compounds. As shown in Fig. 4A, *smeV* gene expression is increased by 764, 4402 and 8972-fold in the presence of clioquinol, iodoacetate and sodium selenite, respectively, in comparison with the level given by the efflux pump basal expression. The same effect is observed in Fig. 4B, where *smeY* gene expression is increased by 6, 47, 71 and 102-fold in the presence of boric acid, chloramphenicol, erythromycin and lincomycin, respectively, when comparing with the *smeY* basal expression level. The obtained results are in agreement with the previous YFP-based obtained results, validating once again the PBT02 sensor strain as well as the new developed PBT10 strain. These data allow us to confirm that the selected compounds from Biolog plates clioquinol, iodoacetate and sodium selenite are *smeVWX* inducers, and boric acid and the antibiotics chloramphenicol, erythromycin and lincomycin are able to trigger *smeYZ* expression.

**Figure 4.**
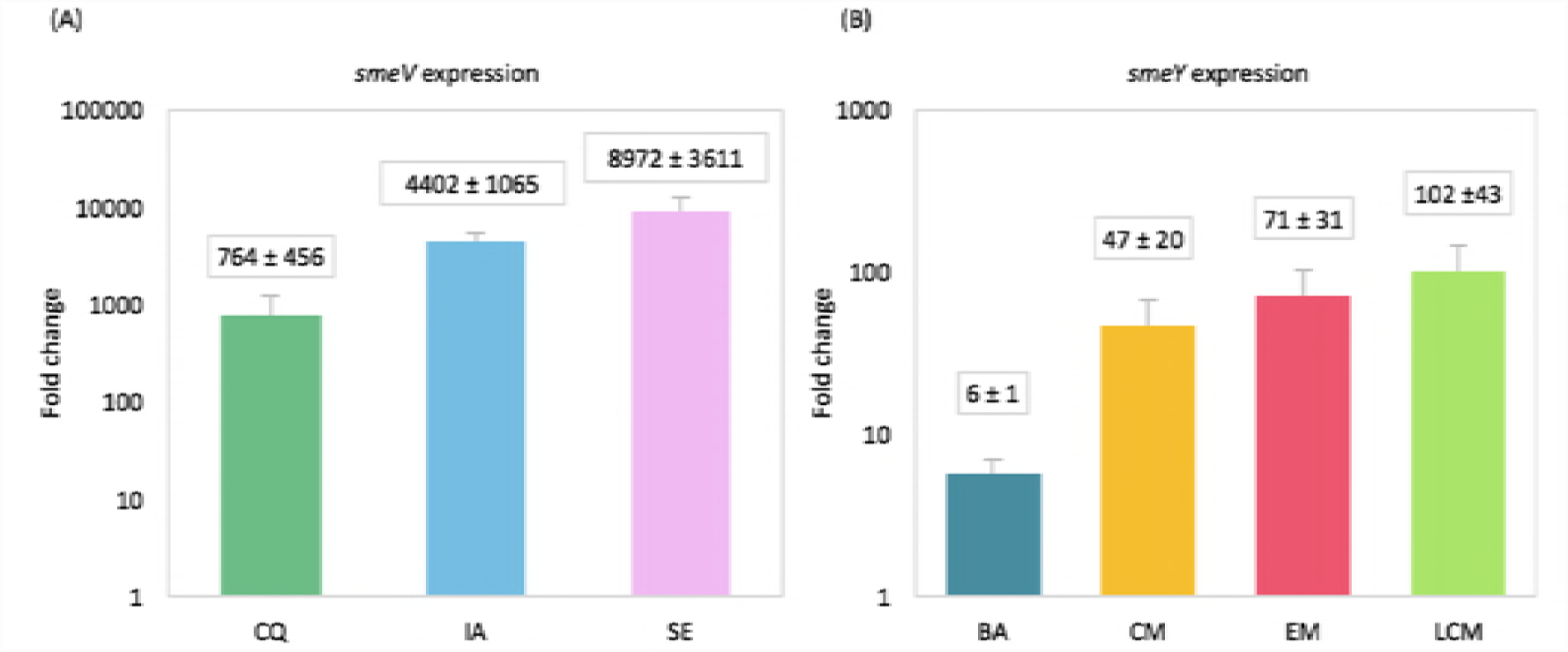
Effect of putative inducers on the mRNA levels of *smeV* and *smeY*. Expresion of *smeV* and *smeY* was measured in the presence and in the absence of their potential inducers by real time RT-PCR. (A) *smeV* expression in D457 strain after 90 min incubation with clioquinol (78 µM), iodoacetate (0.5 mM) or sodium selenite (0.5 mM). (B) *smeY* expression in D457 strain after 90 min incubation with boric acid (6.25 mM), chloramphenicol (1 µg/ml), erythromycin (8 µg/ml) or lincomycin (128 µg/ml). Fold changes were estimated with respect to the values obtained for the untreated D457 strain. Error bars show the standard deviations derived from three independent experiments. CQ, clioquinol; IA, iodoacetate; SE, sodium selenite; BA, boric acid; CM, chloramphenicol; EM, erythromycin; LCM, lincomycin.

### Deletion of either *smeW* or *smeZ* does not alter the susceptibility of *S. maltophilia* to their corresponding inducers

Since new inducer molecules (all of them with antimicrobial properties) of both SmeVWX and SmeYZ multidrug efflux pumps have been identified, we wanted to elucidate whether these compounds were also substrates of their respective efflux pumps. To test this hypothesis, we used the MBS704 (Δ*smeW*) strain (30) and we constructed a D457-derived mutant with a partial deletion in the *smeZ* gene (PBT100 strain). In order to see whether or not the lack of SmeVWX changed *S. maltophilia* susceptibility to its potential inducers, D457 and MBS704 (Δ*smeW*) strains were grown for 20 h at 37 °C in the presence of clioquinol (78 µM), iodoacetate (0.5 mM) or sodium selenite (0.5 mM) and OD_600_ was recorded every 10 min. The same experiment was performed for *smeYZ* inducers, growing D457 and PBT100 (Δ*smeZ*) strains in the presence of boric acid (6.25 mM), chloramphenicol (1 µg/ml), erythromycin (8 µg/ml) or lincomycin (128 µg/ml). If these compounds are being extruded through the efflux pumps, it is expected that both *smeW* and *smeZ*-defective mutants have a deficiency in their growth due to the toxicity of the different compounds. As shown in Fig. 5, no relevant differences are observed in the growth between the wild-type and the efflux pump-defective strains in the presence of the different inducer compounds at the tested concentrations. It is generally assumed that an efflux pump inducer molecules are also extruded by the efflux system; the efflux pump hence confers resistance to such effectors if they are toxic (33-36). However, the data obtained in this work support that efflux pumps do not always confer resistance to their inducers, maybe because these compounds are not substrates of the efflux pumps they induce.

**Figure 5.**
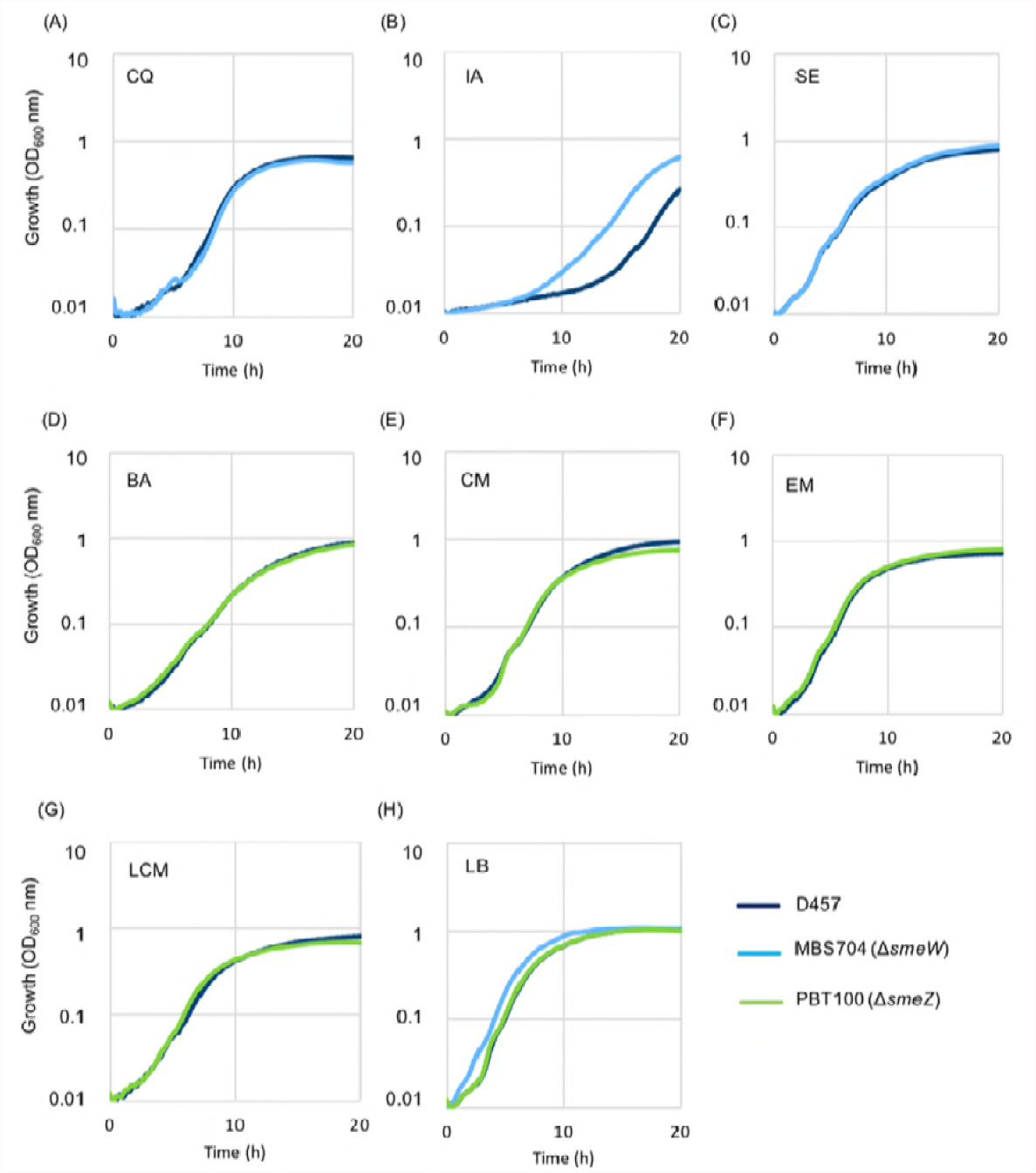
Effect of *smeW* and *smeZ* in the susceptibility of *S. maltophilia* to the inducers of these efflux pumps. The strains D457, MBS704 (Δ*smeW*) and PBT100 (Δ*smeZ*) were grown in LB medium as a control (H). D457 and MBS704 strains were grown in the presence of clioquinol (78 µM) (A), iodoacetate (0.5 mM) (B) and selenite (0.5 mM) (C). D457 and PBT100 strains were grown in the presence of boric acid (6.25 mM) (D), chloramphenicol (1 µg/ml) (E), erythromycin (8 µg/ml) (F) and lincomycin (128 µg/ml) (G). Represented values correspond to the mean calculated from three independent replicates. CQ, clioquinol; IA, iodoacetate; SE, sodium selenite; BA, boric acid; CM, chloramphenicol; EM, erythromycin; LCM, lincomycin.

### The overexpression of MDR efflux pumps promote transient resistance to antibiotics

It is known that overexpression of *smeVWX* contributes to the acquired resistance to chloramphenicol and quinolones of *S. maltophilia* (9), while SmeYZ is able to extrude aminoglycosides, contributing to their intrinsic resistance (10, 11). Since new inducer compounds have been identified for both RND efflux pumps, we wondered whether the susceptibility of *S. maltophilia* would be altered in the presence of these putative effectors, affecting then *S. maltophilia* transient resistance to the known antibiotic substrates of these MDR determinants.

In order to test the contribution of the SmeVWX efflux pump to transient resistance, growth curves were performed in the presence and in the absence of ofloxacin, and with or without sodium selenite, using both the wild-type D457 strain and the Δ*smeW* deletion mutant MBS704. As shown in Fig. 6A and 6B, sodium selenite, at the tested concentration, does not compromise the growth of both strains significantly, whereas ofloxacin impairs the growth of both D457 and MBS704. However, the combination of ofloxacin together with the *smeVWX* inducer sodium selenite does not impede D457 growth. With the aim of assessing whether SmeYZ may contribute to transient resistance when growing in the presence of potential effectors, the same experiment was performed with both D457 and PBT100 (Δ*smeZ*) in the presence or absence of the efflux pump substrate amikacin, and with or without the inducer lincomycin. As shown in Fig. 6C and 6D, lincomycin slightly reduces growth in both strains due to its toxic effect. At the tested concentration of amikacin, the growth is compromised for both strains; however, the presence of lincomycin diminishes the amikacin inhibitory effect. These data suggest that both multidrug efflux pumps are involved in transient resistance to antibiotics when inducers are present, since sodium selenite and lincomycin are able to induce *smeVWX* and *smeYZ*, respectively, changing D457 susceptibility to their antibiotic substrates.

**Figure 6.**
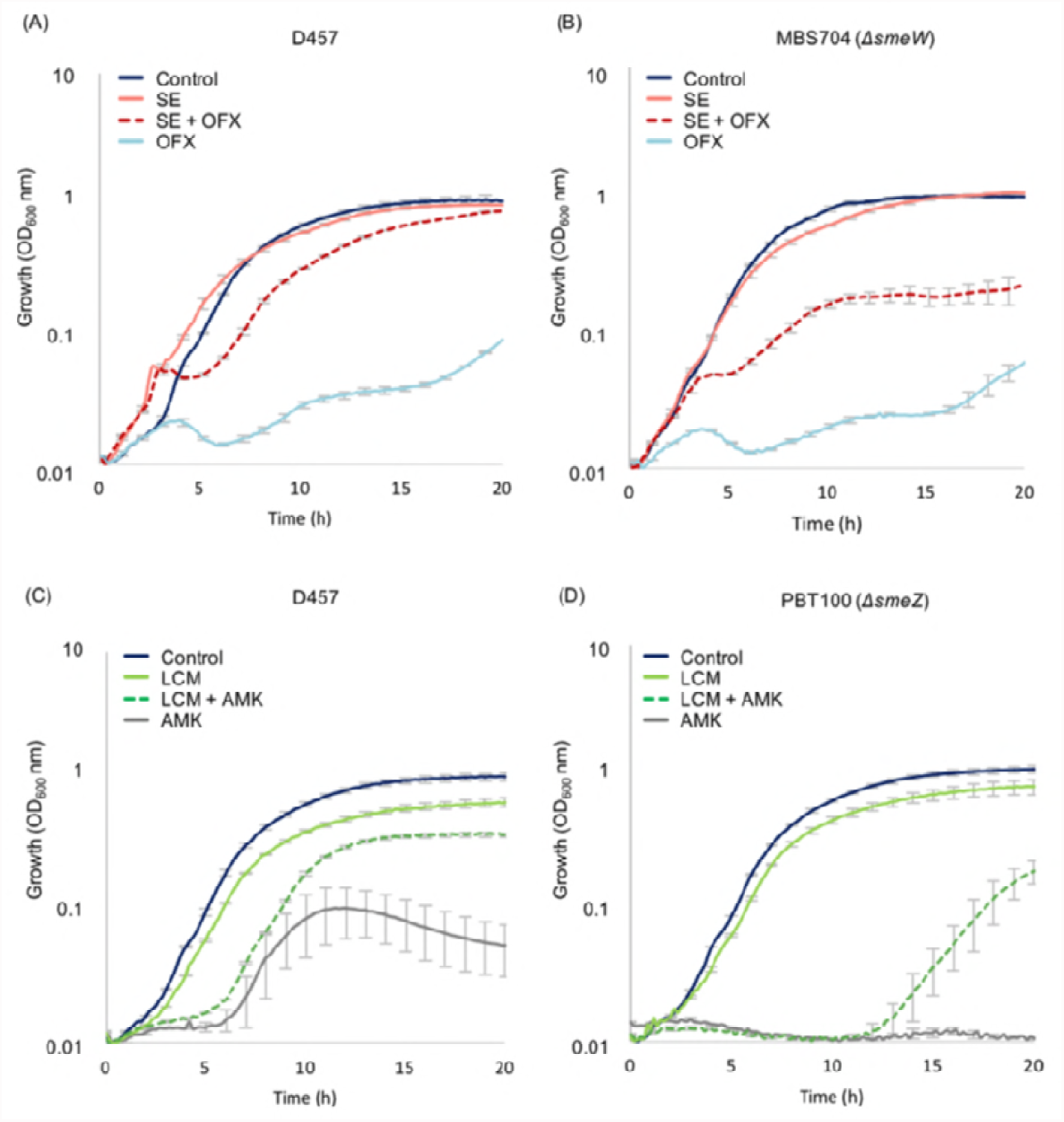
Effect of the inducer compounds sodium selenite and lincomycin in the transient resistance to antibiotics of *S. maltophilia*. Growth curves were performed in the presence of ofloxacin (solid light blue), sodium selenite (solid orange) and both ofloxacin and sodium selenite (dotted red) in D457 (A) and MBS704 (Δ*smeW*) (B) strains. Growth curves were also performed in the presence of amikacin (solid gray), lincomycin (solid light green) and both amikacin and lincomycin (dotted green) in D457 (C) and PBT100 (*ΔsmeZ*) (D) strains. Growth in LB medium is represented in solid dark blue lines for the three strains. Error bars show the standard deviations derived from three independent replicates. SE, sodium selenite; OFX, ofloxacin; LCM, lincomycin; AMK, amikacin.

## Discussion

MDR efflux pumps expression is controlled by transcriptional regulators that act at different levels. In addition to the direct regulation carried out by these regulators, one important aspect to take into account is the overexpression of MDR efflux pumps triggered by the presence of inducer molecules in the medium (13, 37). The conditions that lead to this transient expression of efflux systems, which consequently could give rise to antibiotic resistance induction, are not fully explored and their detection is usually difficult through the use of classical susceptibility methods (13). In this study, we have developed a new methodology for the systematic analysis and identification of MDR efflux pumps’ inducers by combining the Biolog phenotype microarray technology with the use of fluorescent bioreporters.

With the aim of identifying conditions or compounds that would induce the expression of the *S. maltophilia* SmeVWX and SmeYZ efflux pumps, YFP-based reporter strains were developed. In a previous screening using the PBT02 (*P_VWX_*) strain, we identified that vitamin K3, among 30 tested compounds, was an inducer of the SmeVWX system (30). In the current study, two new reporter strains were constructed. The PBT10 strain contains the YFP protein under the control of SmeYZ efflux pump promoter and the PBT03 strain, developed as a positive fluorescence control since harbors the YFP protein under the control of the *EM7* constitutive promoter. Employing Biolog phenotype microarrays, approximately 40 compounds were selected as potential candidates for being *smeVWX* or *smeYZ* inducers.

Among all the potential inducer compounds derived from the Biolog data analysis, clioquinol (5-chloro-7-iodo-8-hydroxyquinoline), iodoacetate, sodium selenite (*smeVWX* inducer candidates), boric acid, chloramphenicol, erythromycin and lincomycin (*smeYZ* inducer candidates), were selected for further analysis. The fluorescence data obtained through the use of Biolog plates were confirmed testing cognate concentrations of the selected agents. The observed induction for both RND efflux pumps was also confirmed by flow cytometry at mid-exponential growth phase after 90 min treatment with the corresponding compounds and by RT-PCR under the same conditions, obtaining similar fold change results in the expression level than those given by the flow cytometry assay. Hence, the Biolog technology has allowed us to successfully identify a set of inducer molecules, being extended for this purpose to other microbial systems.(38)

MDR efflux pumps, besides antibiotic resistance, contribute to different aspects of bacterial physiology, including bacterial interaction with hosts (human, animals and plants) or detoxification of cellular toxic metabolites (39). MDR efflux pumps can also take part of general mechanisms of response to cellular stress that contribute to ameliorate the adverse effects caused by stressors agents or stress by-products (e.g. oxidative stress or envelope stress) (18, 40). We hypothesize that when MDR efflux systems are participating in this global cellular stress response, a general common inducer mechanism of stress could be identified through the analysis of their inducer compounds, despite their structural diversity. As described in this work, clioquinol, iodoacetate and sodium selenite have been identified as *smeVWX* inducers, in addition to the previously described vitamin K_3_ (30). Although clioquinol, sodium selenite and vitamin K_3_ are known to generate oxidative stress (41-44), we determined in our previous work that tert-butyl hydroperoxide, another oxidative stress agent, was not able to induce *smeVWX* (30). In addition, iodoacetate is not related to oxidative stress production (45). However, all of the identified inducer compounds are associated with thiol-reactivity. For instance, iodoacetate is an alkylating reagent that modify thiol groups in proteins by S-carboxymethilation (45); selenite is also known to catalyze the oxidation of thiol groups and to induce protein aggregation (46); clioquinol, as a Cu ionophore, can deliver metal ions into cells, where it exerts its activity through the interaction with thiol and amino groups (47); finally, the previously identified vitamin K3 (menadione) also contributes to redox cycling and has alkylating properties, reacting as well with thiol groups (44). All of these evidences suggest that the *smeVWX* induction mechanism might be related, at least in part, to the thiol-reactivity of the inducer compounds.

In the case of the SmeYZ efflux pump, boric acid, erythromycin, chloramphenicol and lincomycin were identified as inducers. Erythromycin, a macrolide antibiotic, targets the 50S subunit of the bacterial ribosome and inhibits the nascent chain elongation (48). Chloramphenicol is another antibiotic whose target is the 50S subunit of the ribosome by its binding to the peptidyl-transferase center (PTC), where peptide-bond formations happen (49). Both erythromycin and chloramphenicol are also known to directly inhibit the 50S subunit biogenesis (50). The third identified antibiotic inducer, lincomycin, a lincosamide family antibiotic, also targets the 50S subunit of the ribosome by inhibiting the peptide-bond formation, as chloramphenicol (51). *smeYZ* was also found to be induced by boric acid, which is not an antibiotic but impairs the acylation of tRNAs and inhibits protein synthesis (52). All of these compounds share a common mechanism of action, suggesting that SmeYZ mechanism of induction might be related to protein synthesis inhibition. Supporting this possibility, other compounds from Biolog plates, such as oleandomycin, spiramycin, tylosin, penimepiclyne or fusidic acid, whose mechanism of action is also protein synthesis inhibition, were found to increase the YFP levels given by the PBT10 reporter strain (Table 2). These data reinforce the hypothesis that ribosome-stalling stress could be a signal that triggers *smeYZ* expression. It has been previously reported that ribosome-targeting antibiotics are able to induce the expression of the *Pseudomonas aeruginosa* MexXY efflux pump through the induction of the expression of the PA5471 gene (53). In our case, it is possible that SmeYZ plays a role in exporting anomalous peptides or processed by-products alleviating ribosome-stalling stress. Notwithstanding these evidences, the molecular basis underlying the mechanism remain to be established.

It has been described that RND efflux pumps extrude several of their known inducers, such as aminoglycosides by MexXY-OprM in *P. aeruginosa* (54), or triclosan by SmeDEF in *S. maltophilia* (33). The extrusion of the inducer molecules would be a protection mechanism against these toxic agents. However, in this work we have described that the lack of either SmeVWX and SmeYZ does not affect *S. maltophilia* growth from the action of their toxic inducers (Figure 5). Different circumstances have been proposed to explain this apparent contradiction: one possibility is that the concentration for induction is lower or similar to the toxic concentration of the tested compound, not allowing to detect an effect (26). Also, the bacterium can contain more efficient mechanism, (e. g. other efflux pumps) able to extrude or detoxify the same compounds, needing to be removed for the observation of an effect in the bacterial growth (55). The possibility that an effector is not extruded by the efflux pump it induces cannot be disregarded. In this respect, it is worth to notice that both RND substrates and inducers of a given efflux pump are frequently structurally very diverse (39), although they can interfere with the same cell machinery or target. Indeed, we have identified in this work two common mechanisms of stress generated by a set of RND inducers, each one inducing the expression of an efflux pump. It is important to consider that RND efflux pumps mainly extrude substrates that are located in the periplasm (56), while the regulation of these systems take place in the cytoplasm (57). This differentiated compartmentalization may justify the possibility that RND inducers might not always be substrates of efflux pumps.

This means that in some situations, MDR efflux pumps might be overexpressed despite there is no apparent advantage for the bacteria. However, the presence of the inducer may imply a situation of transient resistance to other toxic compounds which represents a benefit in some conditions. We have assessed this situation for clinically used antibiotics that are RND efflux pumps substrates using sodium selenite and lincomycin, the strongest inducers of SmeVWX and SmeYZ efflux pumps, respectively. Both inducer molecules were able to promote transient reduced susceptibility to ofloxacin (SmeVWX substrate) or amikacin (SmeYZ substrate).

This is a fact to take into account regarding clinical situations, since some of these inducer molecules could be provided during a *S. maltophilia* infection treatment. For instance, sodium selenite has been recently administered during a phase I clinical trial in terminal cancer patients (58) due to its cytotoxic effect on proliferating cancer cells. Since these patients are delicately vulnerable to infections caused by MDR Gram-negative bacteria, such as *S. maltophilia* (59), the dispensation of sodium selenite could lead to the overexpression of *smeVWX* and thus, transient resistance to its substrates. The clinical consequences of this situation need to be further investigated.

## Conclusions

Through the combination of Biolog phenotype microarrays and fluorescence-based reporter strains, we have developed a new high-throughput methodology to identify MDR efflux pumps inducer compounds. The common mechanism of action of the detected inducer molecules has allowed us to establish possible mechanisms of induction of both *smeVWX* and *smeYZ* in *S. maltophilia*. *smeVWX* is likely induced by those compounds related to thiol-reactivity, while *smeYZ* is induced by those agents that target the ribosome, suggesting a relationship between the expression of the efflux pump and the inhibition of protein synthesis. These results indicate that MDR efflux pump expression is not always triggered by a specific compound, but by stress signals generated by certain molecules with the same mechanism of action.

The knowledge of these environmental circumstances and/or the inducer molecules, is of importance in terms of finding these agents *in vivo*, where they could compromise an antibiotic therapy due to the induction of MDR efflux pumps (40). In our work, we have determined that the presence of sodium selenite provides with ofloxacin resistance due to the presence of SmeVWX; also, lincomycin is able to lead to amikacin resistance because of the overexpression of *smeYZ*.

## Materials and Methods

### Bacterial strains and growth conditions

All plasmids and strains derived from this study are listed in Table 1. All experiments were performed using LB medium at 37 °C. The following antibiotics were added when required: streptomycin (Sm; 50 µg/ml) for *E. coli* harboring the pSEVA4413 plasmid, ampicillin (Ap; 100 µg/ml) for *E. coli* containing the pGEM-T Easy vector and the pGEM-T-derived plasmids pPBT02 and pPBT11, and kanamycin (Km; 50 µg/ml and 500 µg/ml) for *E. coli* and *S. maltophilia*, respectively, for the selection of pSEVA237Y and the derived plasmids pPBT04, pPBT05 and pPBT08. Medium was supplemented with 0.5 mM isopropyl-β-D-thiogalactopyranoside (IPTG) and 80 µg/ml 5-bromo-4-chloro-3-indolyl-β-D-galactopyranoside (X-Gal) for the induction and detection of β-galactosidase production.

### Plasmid constructions

The PBT10 strain, which harbors the reporter plasmid pPBT08 containing the *smeYZ* promoter region, was obtained as described previously (30) using the primers SmeYZ_F (5’-<u>GAATTC</u>GGCGGGGCCGTAAG-3’, EcoRI site underlined) and SmeYZ_R (5’-<u>AAGCTT</u>TGCTGTGCACAATG-3’, HindIII site underlined). In order to obtain the PBT03 strain, which harbors the pPBT05 plasmid with the constitutive promoter *EM7*, pSEVA4413 plasmid was digested with PacI and HindIII restriction enzymes (New England BioLabs). The product corresponding to the *EM7* promoter was purified from a 1 % agarose gel using a DNA purification kit (GE Healthcare) and cloned into pSEVA237Y plasmid, previously digested with the same restriction enzymes, using T4 DNA ligase (New England BioLabs). The resulting plasmid pPBT05 was introduced by transformation in *E. coli* CC118*λpir*. Then, pPBT05 was introduced into *S. maltophilia* D457 by tripartite matting as previously described (30).

### Deletion of *smeZ*

A partial deletion of the *smeZ* gene was performed in *S. maltophilia* D457 through homologous recombination. 545-bp (ZA) and a 549-bp (ZB) fragments from the 5’ end and 3’ end of the *smeZ* gene, respectively, were amplified by PCR using primers ZAF (5’-<u>GAATTC</u>ATGGCACGTTTCTTCATCGATCGCCCGGTGTTCGC-3’, EcoRI site underlined) and ZAR (5’-ATCGACAACAACAGCAGCCATGCTCGGCACCGAACAACTG-3’), and ZBF (5’-CAGTTGTTCGGTGCCGAGCATGGCTGCTGTTGTTGTCGAT-3’) and ZBR (5’-<u>GAATTC</u>TCAACGATGTTCCGTTCCATCCACGGTTCCTCCCGGC-3’, EcoRI site underlined). An overlapping PCR was performed using ZA and ZB fragments as template using ZAF and ZBR primers, obtaining a 1000-bp fragment (ZAB). This ZAB fragment was purified from a 1 % agarose gel using a DNA purification kit (GE Healthcare) and cloned into pGEM-T Easy (Promega) following the manufacturer’s protocol. The sequence was confirmed by DNA sequencing. Afterwards, this plasmid was digested using EcoRI (New England BioLabs) and the resulting ZAB fragment was cloned into the suicide vector pEX18Tc (60), obtaining the pZAB7 plasmid, which was introduced by transformation into CC118*λpir*. Selection was performed using tetracycline (4 μg/ml). pZAB7 was introduced by tripartite matting into *S. maltophilia* D457 (61) and selection was performed on LB agar plates with tetracycline (12 μg/ml) and imipenem (20 μg/ml). Tet^R^ colonies were streaked onto 12 μg/ml tetracycline plates and 10 % sucrose plates. Tet^R^ and Sac^S^ colonies were streaked onto 10 % sucrose plates and incubated at 30 °C overnight. From the sucrose plates, Sac^R^ colonies were streaked onto 12 μg/ml tetracycline plates and 10 % sucrose in order to obtain double recombinants with the partial deletion of *smeZ* gene. Deletion was confirmed in the PBT100 strain using primers FragZAB_L (5’-GTGCAGAACCGGATCAAG-3’) and FragZAB_R (5’-CGAACTCGACAATGAGGAT-3’), and InternoSmeZ_F (5’-CGGTGTCGATCCTGTTCT-3’) and InternoSmeZ_R (5’-TGGATCGAGGTCATGAAATA-3’).

### Biolog phenotype microarray assay

The phenotypic microarray experiment was performed using the 96-well plates PM11-16 from the Biolog Phenotypic Microarray (PM) System (Biolog), which contain a variety of chemical agents including antimicrobials (62). All tested compounds are listed in Table S1. Each compound is found in four different quantities in the plates. To carry out the experiment, 100 μl of LB medium were added to each well and plates were incubated at room temperature under constant shaking for 2 h. The purpose of this first shaking incubation is to dissolve the different compounds in the medium before inoculating the bacterial cells, since the different chemical agents are dried to the base of each well. Ten microliters of an overnight cell culture were added to each well to a final OD_600_ of 0.01. Plates were then incubated at 37 °C for 20 h using a Tecan Spark 10M plate reader (Tecan), measuring growth (OD_600_) and fluorescence every 10 min. Plates were shaken for 5 s every 10 min before each measurement. For yellow fluorescent protein (YFP) detection, the plate reader was set with an excitation wavelength at 508 nm and emission wavelength at 540 nm.

### Normalization of the results

The analysis of the promoter activity for each reporter strain was performed following a two-step method. First, fluorescence curves were fitted to a first order differential equation, analogously to the one that represents bacterial growth. In this way, the maximum specific rate of fluorescence production (*µ*) was calculated according to the equation *F (t) = F_0_ e^µ t^* [being F (t): fluorescence across the time; F_0_: initial fluorescence; *µ*: maximum specific rate of fluorescence production (h^−1^) and t: time (h)]. The parameter was obtained for the *EM7* (*µ_EM7_*), *smeYZ* (*µ_YZ_*) and *smeVWX* (*µ_VWX_*) promoters for each specific compound and concentration. Secondly, a normalization process was done with the *µ_YZ_* and *µ_VWX_* by dividing them by the value of *µ_EM7_* for each compound and concentration. We stablished a threshold of 1.5 x IQR (interquartile range) of each data set including all the compounds and concentrations where detectable growth was observed to define overexpression.

### Confirmation of the induction of gene expression

Sodium selenite, clioquinol (5-chloro-7-iodo-8-hydroxyquinoline) and iodoacetate were selected as possible inducers of the SmeVWX efflux pump, while boric acid and the antibiotics erythromycin, chloramphenicol and lincomycin were chosen as possible inducers of the SmeYZ efflux pump. First, minimum inhibitory concentration (MIC) was determined for each compound by the microdilution method in LB using 96-well plates (NUNCLON^TM^ Δ Surface). Ten microliters of an overnight cell culture were added to 140 μl of medium containing different concentrations of the selected compound to a final OD_600_ of 0.01. Plates were incubated at 37 °C for 20 h without shaking. MIC value was defined by the lowest concentration at which bacterial growth was not observed in the tested compound.

Three concentrations of each compound were chosen for the induction assay considering the obtained MIC values: sodium selenite (0.25, 0.5, and 1 mM), clioquinol (78, 156, and 312 µM), iodoacetate (0.25, 0.5, and 1 mM), boric acid (3.125, 6.25, and 12.5 mM), erythromycin (4, 8, and 16 µg/ml), chloramphenicol (0.5, 1, and 2 µg/ml) and lincomycin (64, 128, and 256 µg/ml). Ten microliters of the bacterial culture were added to 140 µl of medium to a final OD_600_ of 0.01 using Corning Costar 96-well black clear-bottom plates (Corning Incorporated). Plates were incubated at 37 °C for 20 h, and growth (OD_600_) and fluorescence were monitored every 10 min in the plate reader Tecan Spark 10M (Tecan). Plates were shaken for 5 s every 10 min before each measurement. An excitation wavelength at 508 nm and emission wavelength at 540 nm were set for YFP detection.

### Flow cytometry assays

*S. maltophilia* reporter strains PBT02 (*P_VWx_*), PBT03 (*P_EM7_*) and PBT10 (*P_YZ_*) were inoculated in 100 ml Erlenmeyer flasks with 20 ml of LB medium to a final OD_600_ of 0.01 and incubated at 37 °C with shaking. When bacterial cultures reached mid-exponential growth phase (OD_600_ ≈ 0.6), induction with the different compounds was performed using the following concentrations: 0.5 mM sodium selenite, 78 µM clioquinol, 0.5 mM iodoacetate, 6.25 mM boric acid, 8 µg/ml erythromycin, 1 µg/ml chloramphenicol, and 128 µg/ml lincomycin. A bacterial culture with no compound was used as control. Cells were then incubated with shaking for 90 min at 37 °C. One milliliter of each culture was centrifuged at 13.000 rpm for 1 min at room temperature. Cells were washed with 500 µl of PBS and centrifuged as indicated above. The bacterial pellets were suspended with 300 µl of 0.4 % paraformaldehyde and incubated at room temperature for 10 min. Cells were then centrifuged as above indicated and suspended in 1 ml of PBS. In order to avoid false positive signals, all the media and buffers employed were filtered through 0.22 µm pore filters (Corning Incorporated). With the aim of measuring YFP production at the single-cell level, samples of each reporter strain containing 20.000 cells were analyzed using the Gallios^TM^ flow cytometer (Beckman Coulter). Data processing was accomplished with Kaluza 1.5 software (Beckman Coulter).

### RNA preparation and real-time RT-PCR

RNA was obtained as described previously with some modifications (30). Briefly, a 100 ml Erlenmeyer flask with 20 ml of LB medium was inoculated with an overnight culture of *S. maltophilia* D457 to reach a final OD_600_ of 0.01. Cell cultures were incubated at 37 °C in agitation until they reached mid-exponential growth phase (OD_600_ ≈ 0.6). At this point, the induction assay was performed adding the different compounds at the concentrations required: 0.5 mM sodium selenite, 78 µM clioquinol, 0.5 mM iodoacetate, 6.25 mM boric acid, 8 µg/ml erythromycin, 1 µg/ml chloramphenicol, and 128 µg/ml lincomycin. A bacterial culture with no compound was used as control. Cultures were incubated with shaking for 90 min. Ten milliliters of each culture were taken and centrifuged at 8.000 rpm for 20 min at 4 °C. RNA extraction was performed as described (30) and cDNA was obtained using the High-Capacity cDNA reverse transcription kit (Applied Biosystems). Hundred nanograms of cDNA were used for each reaction. Real-time RT-PCR experiment was performed in an ABI Prism 7500 real-time system (Applied Biosystem) using the Power SYBR green PCR master mix (Applied Biosystem). Primers RT-SmeV.L (5’-GTCGACTTCCTCGACAACC-3’) and RT-SmeV.R (5’-TTGCCATCCTTGTCTACCAC3’) were used to amplify *smeV*; primers RT-SmeY.L (5’-CATTGGTGACCGAAGGTG-3’) and RT-SmeY.R (5’-TTGATACCGGAGAACAGCAG-3’) were used to amplify *smeY*; and primers RT-ftsZ.L (5’-ATGGTCAACTCGGCAGTG-3’) and RT-ftsZ.R (5’-CGGTGATGAACACCATGTC-3’) were used to amplify the housekeeping gene *ftsZ*. Relative changes in gene expression were determined according to the 2^-ΔΔ*CT*^ method (63). Mean values were obtained from three independent replicates in each experiment.

### Determination of transient resistance to antibiotics

In order to determine the contribution of SmeVWX and SmeYZ to transient resistance to antibiotics, the respective inducer molecules of each efflux pump, sodium selenite and lincomycin, were selected to carry out growth curves in the presence of antibiotics. D457 and MBS704 (Δ*smeW*) strains were grown in ofloxacin (2 µg/ml), sodium selenite (0.5 mM), or ofloxacin and sodium selenite in combination at the same concentrations. On its hand, D457 and PBT100 (Δ*smeZ*) strains were grown in the presence of amikacin (16 µg/ml), lincomycin (128 µg/ml), or amikacin and lincomycin in combination at the same concentrations. Ten microliters of overnight cultures of the three *S. maltophilia* strains were added to 140 µl of LB medium containing the different compounds to a final OD_600_ of 0.01 in 96-well plates (NUNCLON^TM^ Delta Surface). Plates were incubated at 37 °C for 20 h using a Tecan Spark 10M plate reader (Tecan) and growth (OD_600_) was recorded every 10 min. Plates were shaken for 5 s before every measurement.

## Authors’ contributions

PB. performed the experiments and contributed to the design of the study. FC contributed to the experiments and the design of the study. JLM contributed to the design of the study. All authors contributed to the interpretation of the results and to the writing of the article.

## Supplemental material

**Table S1.**
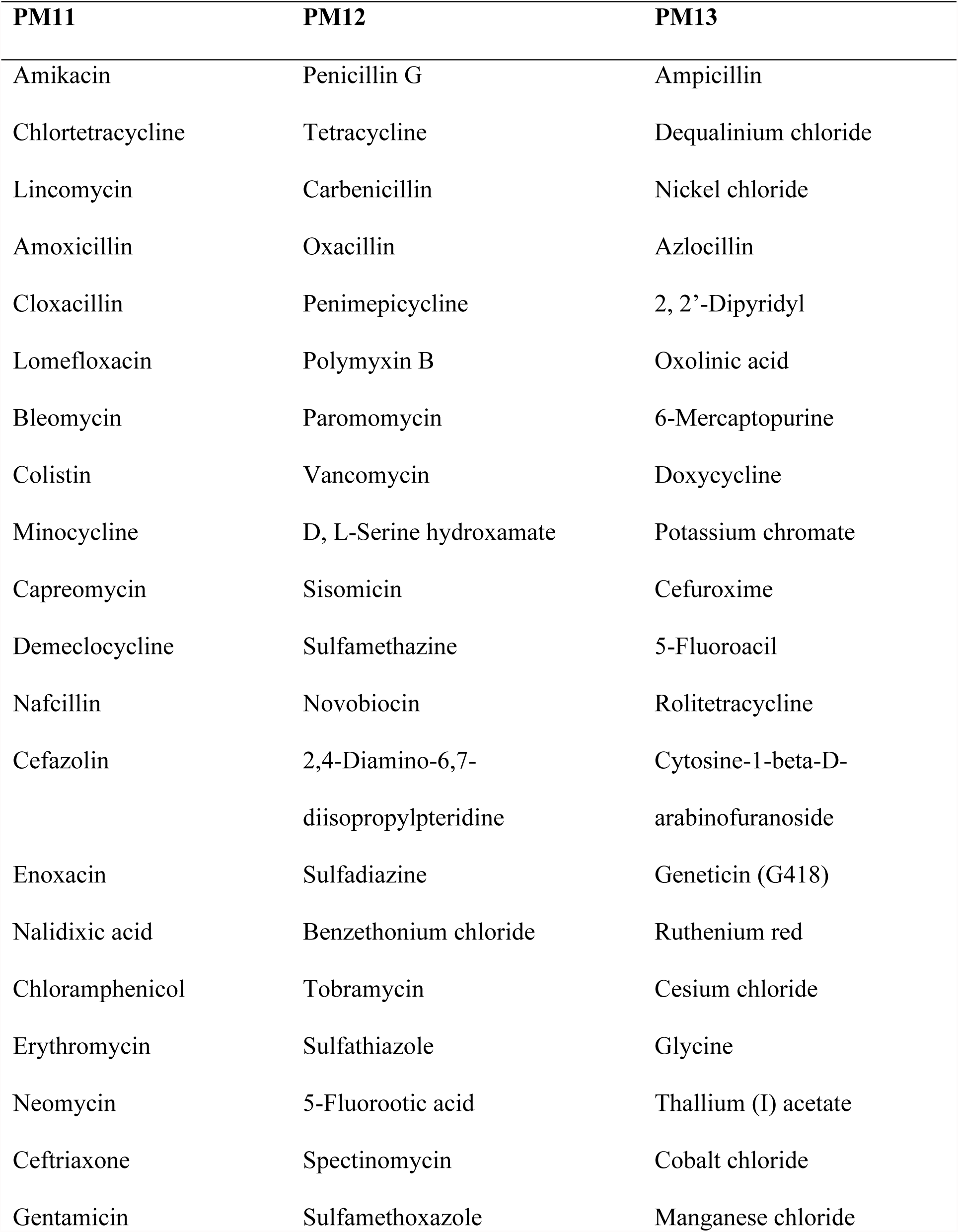
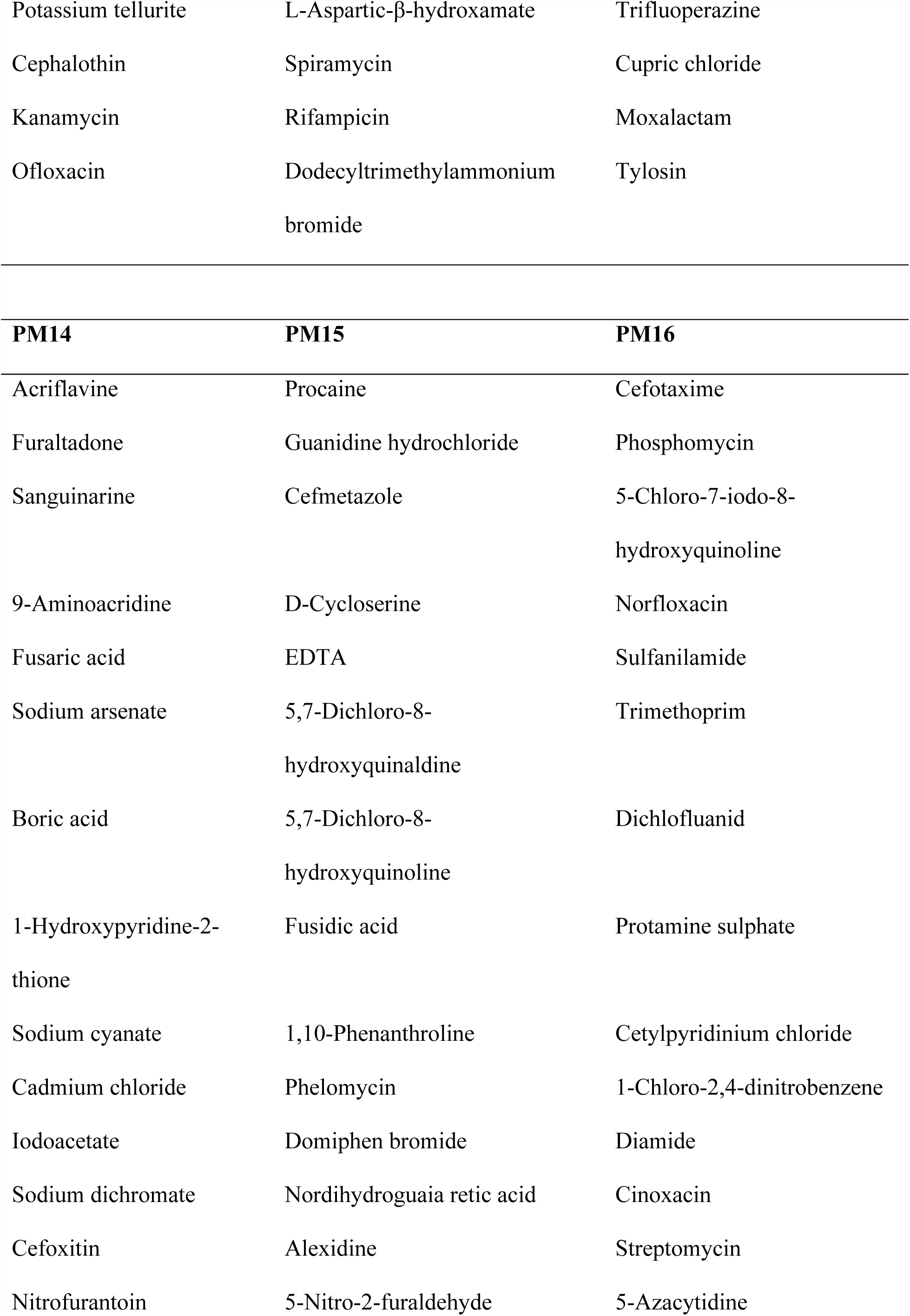
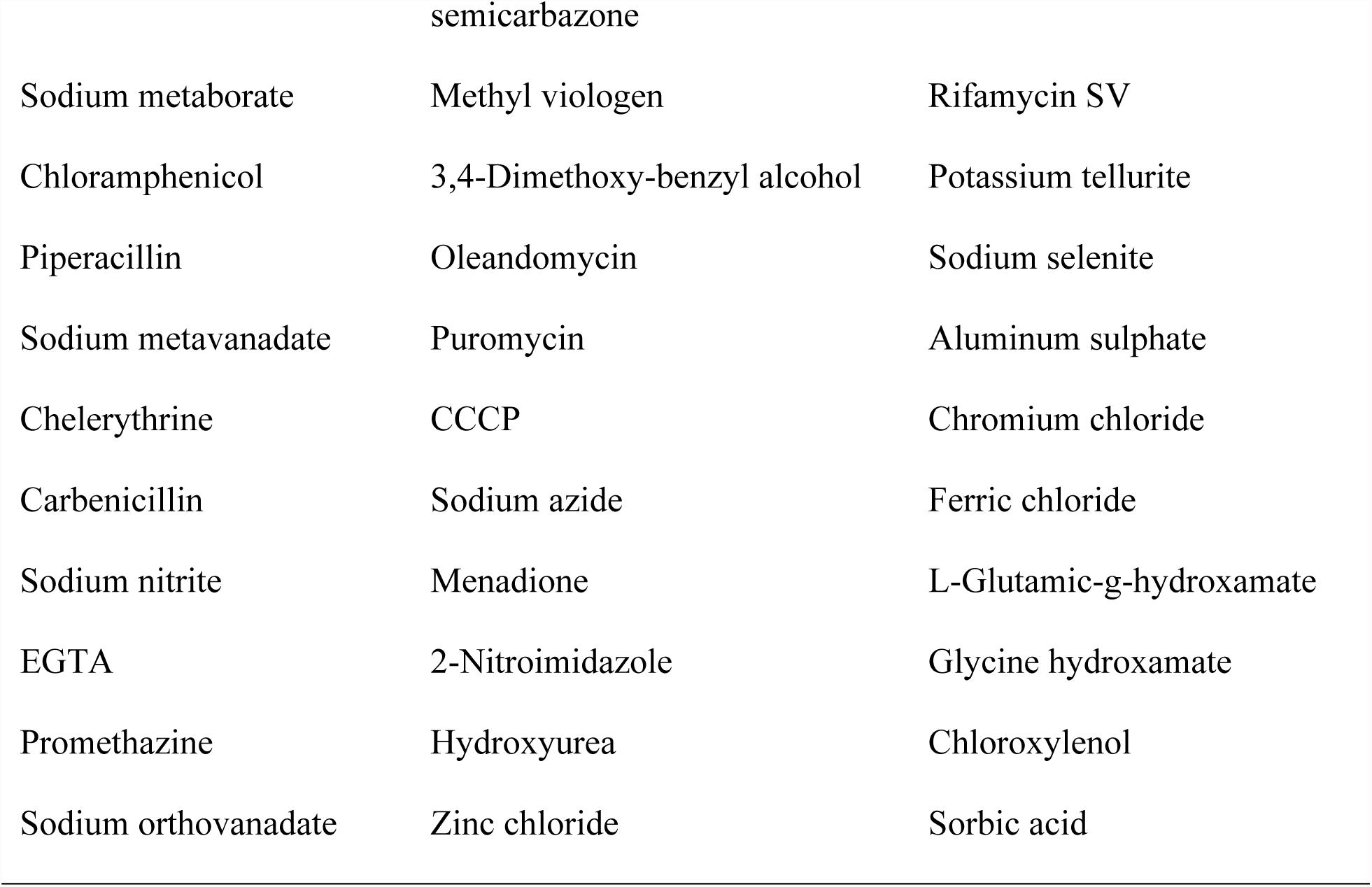
Biolog phenotype microarray tested compounds

**Figure S1.**
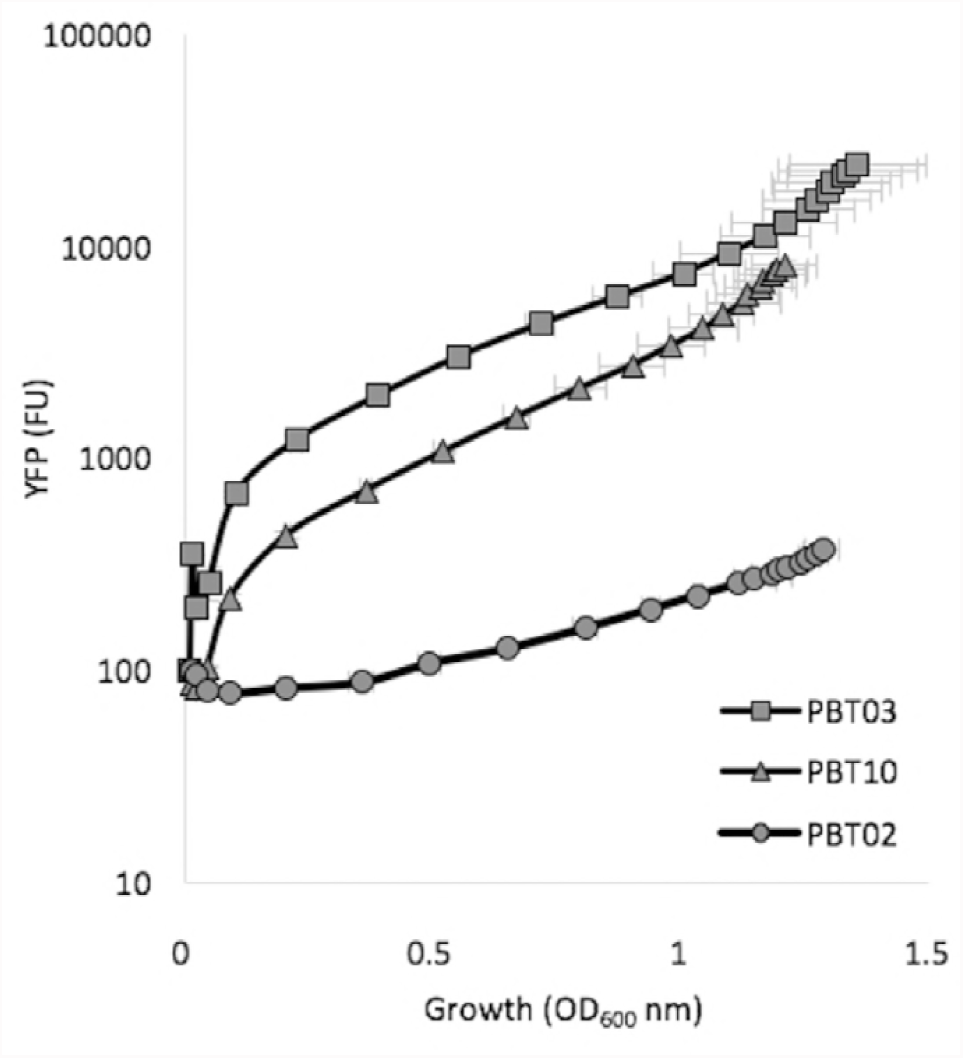
Growth and fluorescence values given by sensor strains PBT03 (*P_EM7_*), PBT10 (*P_YZ_*) and PBT02 (*P_VWx_*). Error bars show standard deviations from three independent replicates. FU, fluorescence units.

**Figure S2.**
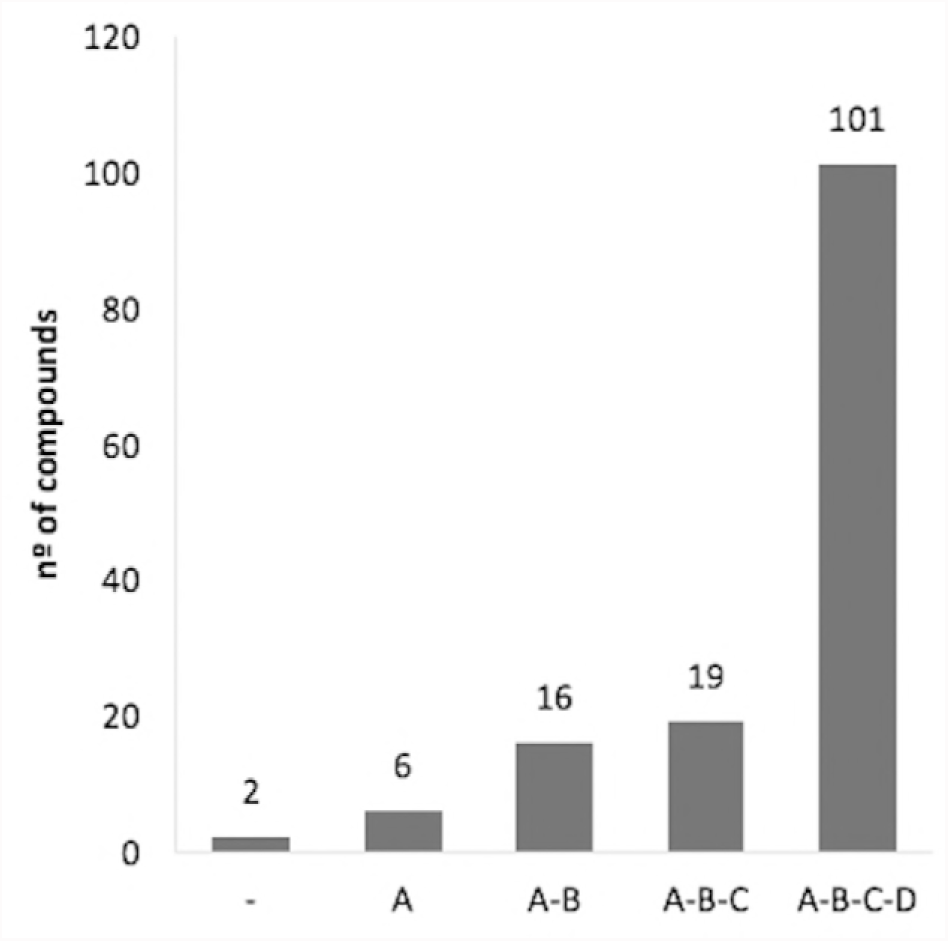
Compounds concentrations where *S. maltophilia* growth is detected. Compounds are divided in five categories. A-B-C-D compounds represent those agents where *S. maltophilia* growth is observable in all concentrations. A-B-C compounds are those where *S. maltophilia* grows only in A, B and C concentrations. The same reasoning for A-B and A compounds. No growth was observed for – compounds.

